# Evaluating the Functional Pore Size of Chloroplast TOC and TIC Protein Translocons

**DOI:** 10.1101/188052

**Authors:** Iniyan Ganesan, Lan-Xin Shi, Mathias Labs, Steven M. Theg

## Abstract

The degree of residual structure retained by proteins while passing through biological membranes is a fundamental mechanistic question of protein translocation. Proteins are generally thought to be unfolded while transported through canonical proteinaceous translocons, which has historically been the thought for the translocons of the outer and inner chloroplast envelope membranes (TOC and TIC). Here, we readdressed the issue and found that medium-sized tightly folded proteins such as the 22 kDa dihydrofolate reductase (DHFR) can be tolerated by TOC and TIC. Chimeric DHFR fused with RuBisCO small subunit transit peptide (tp22DHFR) was found to be imported into chloroplasts in complex with its stabilizing ligand, methotrexate (MTX), in a folded conformation. Following import, both mature tp22DHFR and MTX were found in the chloroplast stroma. A subsaturating concentration of MTX was used to exclude the possibility that MTX was stripped off tp22DHFR, independently imported into the chloroplasts, and reassociated with imported tp22DHFR. Independent MTX import was further excluded by use of fluorescein conjugated MTX (FMTX), which has very slow membrane transport rates relative to unconjugated MTX. The TOC/TIC pore size was determined by probing the translocons with particles of fixed diameter and found to be greater than 25.6 Å, large enough to support folded DHFR import. The pore size is also larger than those of the mitochondrial protein translocons that have a requirement for protein unfolding.

**SIGNIFICANCE:** The chloroplast TOC and TIC translocons are responsible for the import of up to 95% of all chloroplast proteins and are therefore essential for plastid biogenesis and photosynthesis. However, the mechanisms of protein import into chloroplasts are not well understood. The TOC/TIC translocons have long been suggested to have a strong unfoldase activity relative to other comparable protein translocons. Here, we present data suggesting that this is not true, and that instead, they possess a relatively large pore size. This identifies TOC and TIC as rather unique protein translocons capable of transporting folded proteins across a double membrane barrier, which has important implications in the mechanisms of TOC/TIC function and biogenesis of photosynthetic proteins.

Classification - Biochemistry

## INTRODUCTION

Protein import into chloroplasts is a vital prerequisite for photosynthesis and therefore for plant growth and development. The protein translocons of the outer and inner chloroplast membranes (TOC and TIC, respectively) are responsible for the import of approximately 95% of all chloroplast proteins from the cytoplasm and are highly conserved amongst all land plants (1). The TOC complex is composed of Toc75, Toc34, and Toc159 at a 4:4:1 or 3:3:1 ratio (2). Toc75, a β-barrel BamA ortholog, is the major pore-forming subunit at the outer membrane. The composition of the TIC complex has been disputed, with Tic20, Tic21, and/or Tic110 proposed as major pore-forming subunits atthe inner membrane (2, 3). Structures are available for many of the soluble domains of TOC/TIC components, but little is known about the assembled complex structures. The pore sizes of Toc75, Tic20, and Tic110 have been respectively estimated to be 14-26 Å, 7.8-14.1 Å, and 15-34 Å by electrophysiological measurements in proteoliposomes (4-6). However, these calculations rely on many assumptions and may not reflect the full functional pore size range of the subunits within their native complex and membrane environments. The driving force for protein import derives from ATPase activity of stromal chaperones that exert a pulling force on the N-terminus of precursor proteins. Stromal Hsp70 plays a major role, presumably analogous to the mechanism of mitochondrial matrix-localized mtHsp70 (7, 8). Other chaperones proposed to be involved in parallel and/or in series with Hsp70 are stromal Hsp93, stromal Hsp90, and an enigmatic intermembrane space localized Hsp70 (9).

Although most of the TOC/TIC components are known, the mechanisms of protein translocation are not fully understood. A fundamental question in all protein translocation systems is whether the translocon requires substrate proteins to be unfolded, or allows folded proteins to cross the membrane by means of a larger pore. Proteins are generally thought to traverse membranes in an unfolded conformation, as is the case for mitochondrial membrane translocons (TOM and TIM), bacterial SecYEG, and Sec61 in the ER (10-13). However, pathways that transport folded proteins also exist, such as in peroxisomal import and in Tat transport in bacterial and thylakoid membranes (14,15). Unlike mitochondria, chloroplasts are known to tolerate small folded proteins, such as internally cross-linked 6.5 kDa bovine pancreatic trypsin inhibitor, but have been generally considered to unfold larger proteins (15,16). The 22 kDa dihydrofolate reductase (DHFR) is a model protein for folding/unfolding due to interactions with its stabilizing non-covalently bound inhibitor, methotrexate (MTX). MTX binding is a good indicator for DHFR folding because the kinetics of DHFR unfolding match those of DHFR/MTX complex dissociation, implying unfolded DHFR has no affinity for MTX (17). MTX blocked DHFR translocation through the TOM/TIM and SecYEG complexes by preventing DHFR unfolding, but did not block DHFR transport through the Tat pathway (11,13,18). MTX was observed not to block DHFR import into chloroplasts and both DHFR and MTX were found in the stroma (19). In interpreting this experiment, the authors suggested that the TOC/TIC translocons have a strong unfoldase activity capable of stripping MTX away from DHFR. Here, we report that DHFR/MTX is in fact imported into chloroplasts as a folded complex. Additionally, the TOC/TIC pore size was probed and determined to be significantly larger than its functional cognate pore in mitochondrial TOM/TIM.

## RESULTS

### The DHFR/MTX Complex Is Imported into Chloroplasts with Subsaturating [MTX]

The mechanism of DHFR/MTX import into chloroplasts was previously described as separate transport of protein and inhibitor across the envelope membranes, after which they reassociate. The ability of MTX to independently cross the chloroplast envelope membranes complicates the question of whether or not TOC/TIC unfolds the DHFR/MTX complex. However, DHFR/MTX complex import can be differentiated from independent DHFR and MTX import by monitoring the presence of MTX-bound DHFR in the stroma when the import reaction is conducted with a subsaturating MTX concentration. This method was previously used to show DHFR/MTX complex transport through the cpTat pathway (18). *In vitro*-translated DHFR fused to the RuBisCO small subunit transit peptide ([^3^H]-tp22DHFR) and bound to MTX can be detected by protease resistance conferred by MTX binding (Fig. S1). The protease-resistant degradation product is the size of DHFR and smaller than mature [^3^H]-tp22DHFR due to a 22-residue linker between the cleavable transit peptide and DHFR. [^3^H]-tp22DHFR/MTX binding was saturated at MTX concentrations above 4 nM and was negligible below 1 nM (Fig. 1A). [^3^H]-tp22DHFR was incubated with 133 nM MTX to saturate binding, then desalted and diluted into the import reaction to achieve subsaturating MTX concentrations below 1 nM. Desalting reduced MTX concentrations 660-fold, as determined by MTX absorbance at 304 nm (Fig. S2). MTX-pretreated and-desalted [^3^H]-tp22DHFR was more protease resistant after import relative to the negative control without any MTX treatment (Fig. 1B). Since MTX was subsaturating in the import reaction, this indicates that the originally bound MTX remained bound during translocation. MTX-pretreated [^3^H]-tp22DHFR was less protease resistant than the positive controls where MTX was added to lysed chloroplast stroma samples after import. This difference is most likely due to passive [^3^H]-tp22DHFR/MTX complex dissociation under the subsaturating conditions. [^3^H]-tp22DHFR still utilized the TOC/TIC translocons in the presence of MTX since it competed with a native precursor substrate (Fig. S3). An alternate explanation to folded [^3^H]-tp22DHFR/MTX complex import is that folate transporters in the inner membrane with MTX affinity much higher than that of DHFR are able to actively concentrate MTX in the stroma. This level of high affinity MTX transport has not been described for characterized inner membrane folate transporters (20).

**Fig. 1.**
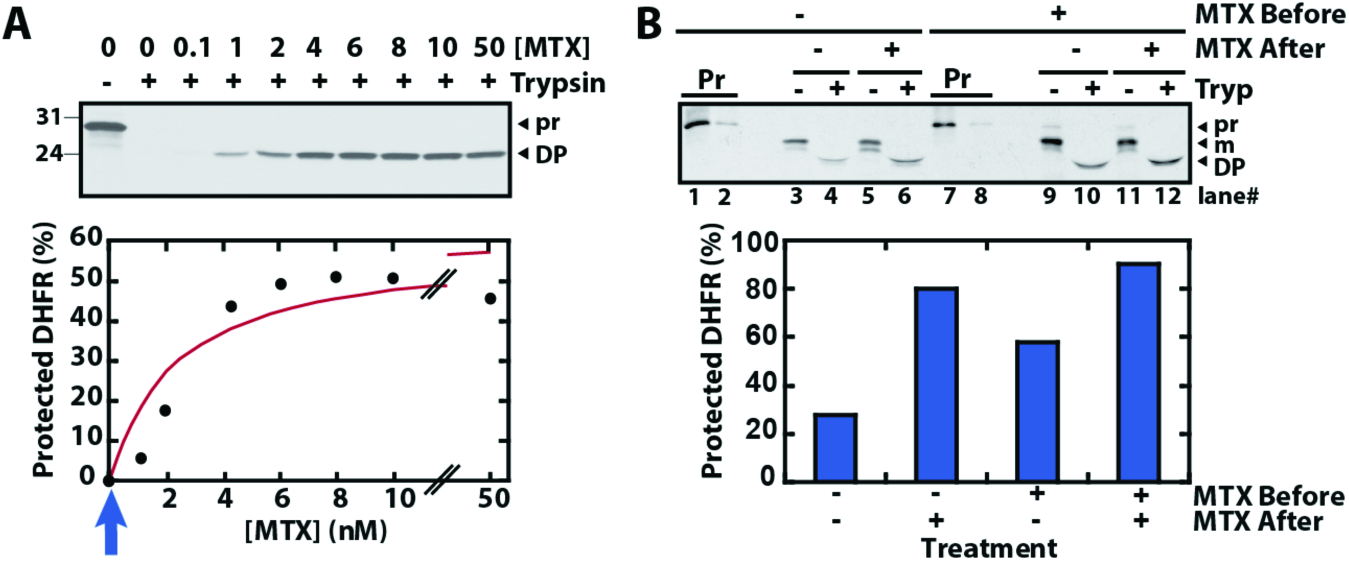
Imported tp22DHFR/MTX complex is detected under subsaturating MTX conditions. (A) Protease protection saturation curve of [^3^H]-tp22DHFR/MTX binding. *In vitro*-translated [^3^H]-tp22DHFR was incubated with methotrexate (MTX) at indicated concentrations on ice for 10 min followed by addition of 200 μg/ml trypsin and 5 min incubation at 37 °C. Reactions were quenched with 2 mg/ml soy bean trypsin inhibitor (SBTI) for 10 min on ice, diluted in Sample Buffer, and analyzed by SDS-PAGE and fluorography. Blue arrow indicates expected MTX concentration of desalted [^3^H]-tp22DHFR/MTX solution used in panel B. **(B)** Import of desalted [^3^H]-tp22DHFR/MTX complex into chloroplasts. Before import, [^3^H]-tp22DHFR was incubated with Import Buffer (IB, Lanes 1-6) or MTX (133 nM, lanes 7-12) on ice for 10 min. The treated precursor samples were then filtered through Zeba Spin Desalting Columns (7K MWCO). Chloroplasts were incubated with above precursor mixtures supplied with 3 mM ATP and 1 mM DTT for 20 min under light. After import, chloroplasts were treated with 200 μg/ml thermolysin on ice for 15 min, washed with IB containing 50 mM EDTA and reisolated by centrifugation. The pellets were resuspended in 15 μl lysis buffer (10 mM MES, 5 mM MgCl_2_, pH 6.5) and sonicated in icy water bath for 30 s. The released stroma was collected by centrifugation at 16000 x *g* for 15 min at 4 °C. MTX (1 μM) or IB was added into stromal samples and the resulting mixtures were then subjected to trypsin (Tryp, 800 μg/ml, lanes 4, 6,10, and 12) or mock (with IB, lanes 3, 5, 9, and 11)) treatments at 37 °C for 10 min. Reactions were quenched by with SBTI (8 mg/mL). Three stromal samples derived from those treated with MTX before import (Lanes 9-12) were combined and TCA precipitated, while lanes 3-6 each derive from a single import reaction. Stromal samples were subjected to SDS-PAGE and fluorography. pr, precursor; m, mature; DP, protease-protected degradation product.

### FMTX Import into Chloroplasts Is Dependent upon DHFR Import

Unlike MTX, fluorescein-conjugated methotrexate (FMTX) has extremely reduced passive membrane diffusion rates (21). This property makes FMTX an ideal substrate to test whether it is imported into chloroplasts independently or in complex with DHFR. FMTX also has enhanced fluorescence when bound to DHFR, which allows for quantification of the DHFR/FMTX complex (22, 23). Such FMTX fluorescence enhancement was seen upon binding chimeric *E. coli* DHFR fused to RuBisCO small subunit transit peptide (tp22EcDHFR) purified from *E. coli*, and was reversed by addition of excess MTX to compete for tp22EcDHFR binding (Fig. 2).

**Fig. 2.**
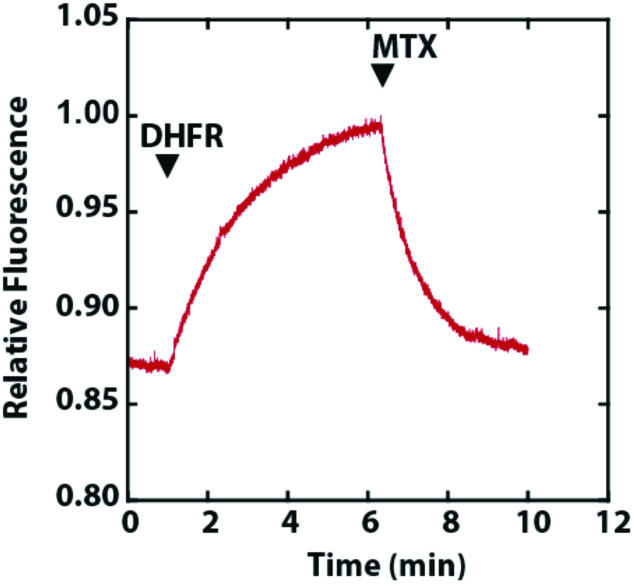
Fluorescence enhancement of FMTX upon binding tp22EcDHFR. tp22EcDHFR (40 nM) was added to 80 nM FMTX followed by addition of excess MTX (800 nM) at indicated time points.

FMTX import into chloroplasts was analyzed by introducing FMTX to chloroplasts during or after tp22EcDHFR import (Fig. 3A). To test for DHFR-dependent FMTX import, both protein and inhibitor were added to the initial import reaction. The reaction was stopped in cold IB and immediately washed to remove excess FMTX. To test for protein-independent FMTX import, tp22EcDHFR was pre-imported so that independently imported FMTX could potentially bind to DHFR in the stroma and accumulate therein. This was achieved by importing tp22EcDHFR in an initial import reaction, followed by washing and thermolysin treatment to remove unimported protein. FMTX was introduced to the chloroplasts in a second mock import reaction with time, temperature, light, and ATP conditions identical to the initial import reaction. The only difference between the DHFR-dependent and-independent FMTX import was the initial location of DHFR (outside or inside the chloroplast, respectively). Thermolysin treatment of chloroplasts should not affect FMTX import since FMTX is small enough to freely pass through pores in the outer membrane and therefore does not require any outer membrane transporters (24).

**Fig. 3.**
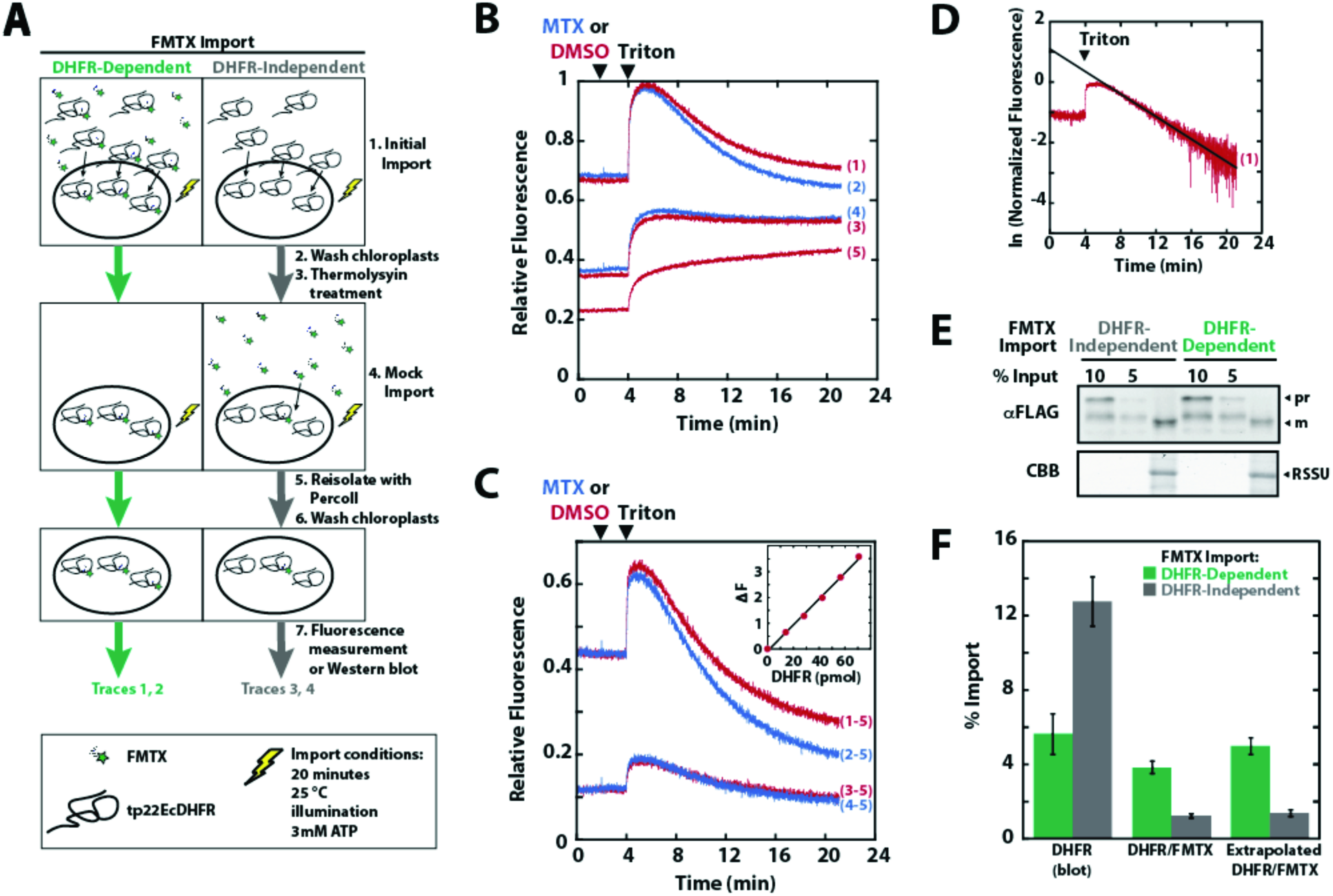
FMTX Import into chloroplasts is dependent upon concurrent DHFR import. **(A)** Schematic of FMTX import experiment. tp22EcDHFR (470 nM) was imported into chloroplasts in 300 μl reactions with or without 17 μM FMTX as shown (Initial Import). Reactions were stopped with cold IB. Chloroplasts were washed and resuspended in 400 μg/mL thermolysin for 1 hour on ice and quenched with 12.5 mM EDTA. Chloroplasts were resuspended in IB with ATP and with or without 17 μM FMTX as shown, and were again exposed to light for 20 min (Mock import). Chloroplasts were reisolated on 40% Percoll cushions, washed with IB, and assayed for fluorescence. **(B)** Fluorescence kinetics of DHFR-dependent FMTX import (traces 1 and 2) or independent FMTX import (traces 3 and 4). Trace 5 represents background chloroplast fluorescence derived from chloroplasts treated as shown in A, except without any addition of protein or FMTX. Excess MTX (17 μM, traces 2 and 4) or DMSO (traces 1, 3, and 5) was added att=2 min and 0.05% Triton-X 100 (TX) was added to all traces at t=4 min. All kinetics represent averages of three independent experiments. **(C)** Trace 5 was subtracted from traces 1 through 4 to remove background fluorescence. Replicates were offset adjusted to the same initial fluorescence. tp22EcDHFR/FMTX complex import was determined as the change in fluorescence due to debinding of FMTX from DHFR (base fluorescence at t=21 min in MTX added trace subtracted from peak fluorescence at t=5 min in the DMSO added trace). (C, Inset) Fluorescence change (ΔF) was calibrated by adding EcDHFR in 14 pmol increments to a saturating 170 nM FMTX solution in IB containing chloroplasts (17 μg/mL Chl). **(D)** The true tp22EcDHFR/FMTX peak at t=4 min was determined by extrapolating linearized DMSO added traces (1 and 3) back to t=4 min. Decay curves were linearized by taking the natural log of normalized data. Sample extrapolation is shown. **(E)** tp22EcDHFR import was determined by α-FLAG blotting, shown along with a Coomassie stained loading control (CBB). **(F)** tp22EcDHFR protein import and tp22EcDHFR/FMTX complex import (non-extrapolated and extrapolated) were quantified from Western blots and fluorescence data, respectively. Error bars indicate SEM (n=3).

All chloroplast samples were reisolated and imported FMTX was detected by fluorescence. The initial fluorescence observed with DHFR-dependent FMTX import was much higher than that of independent FMTX import, supporting the model that the DHFR/FMTX complex was imported (Fig. 3B). Quantification of the imported tp22EcDHFR/FMTX complex requires debinding of FMTX with excess MTX. However, excess MTX did not efficiently replace bound FMTX in the stroma. MTX may not accumulate in the stroma at high enough concentrations to compete for FMTX binding at a detectable rate. Therefore, the chloroplasts were solubilized with 0.05% Triton after addition of excess MTX to allow for complete FMTX debinding. Triton solubilization causes the chloroplast solution to become more transmissive to light, increasing the fluorescence signal at the four-minute time point. Solubilization also causes the effective solution volume surrounding the tp22EcDHFR/FMTX complex to increase 1000-fold from the stromal volume to the total bulk solution volume. This dilution shifts the binding equilibrium towards the unbound state, causing the fluorescence signal to decline at the five-minute time point even without prior addition of excess MTX. This dilution effect is specific to DHFR/FMTX debinding, as it is not seen upon solubilizing chloroplasts with imported precursor covalently conjugated to fluorescein (Fig. S4). In the presence of excess MTX, the fluorescence decay after solubilization was faster due to the additive effects of dilution and MTX/FMTX binding competition (Fig. 3B). The kinetics of FMTX replacement with MTX was seen by subtracting traces with and without MTX addition (Fig. 3B, trace 2-1 and trace 4-3; Fig. S5). This MTX effect is more pronounced for DHFR-dependent FMTX import. To quantify the DHFR/FMTX debinding fluorescence decay, the background chloroplast fluorescence (trace 5) was first subtracted from the other traces (Fig. 3C). The fluorescence change was calibrated by measuring fluorescence of EcDHFR (without tp22 fusion) added stepwise to a solution of chloroplasts and saturating FMTX (Fig. 3C Inset). The fluorescence change was taken as the difference in peak fluorescence without MTX addition (at 5 minutes) and steady state fluorescence after complete FMTX debinding by MTX (at 21 minutes). However, the five-minute peak does not represent the maximum protein-bound FMTX since the dilution effect begins immediately after solubilization at four minutes and competes with the fluorescence increase caused by chloroplast solubilization between four and five minutes. The true peak was therefore determined by linearizing the fluorescence decays and extrapolating back to the four-minute time point when Triton was added (Fig. 3D). Quantitated DHFR-dependent FMTX import was much higher than independent FMTX import, again indicating that folded tp22EcDHFR/FMTX was translocated (Fig. 3F). Replicate samples were analyzed by Western blotto independently quantify tp22EcDHFR import, which matched the extrapolated import of the tp22EcDHFR/FMTX complex (Fig. 3E,F). This convergence of the data suggests a 1:1 ratio of tp22EcDHFR and FMTX import. There was some inhibition of tp22EcDHFR import by FMTX, which suggests that uncomplexed tp22EcDHFR may in fact be somewhat unfolded and thereby imported more efficiently by chloroplasts.

### The TOC/TIC Pore Size Is Greater Than 25.6 Å

To further confirm that folded DHFR can pass through the TOC/TIC translocons, the functional pore size was measured by probing the translocons with particles of fixed diameter attached to precursor proteins. A rigid, spherical, 20 Å monomaleimido Undeccagold particle was covalently conjugated to a single C-terminal cysteine on RuBisCO small subunit containing FLAG and HIS tags for detection and purification (RSSUFHC). The Undeccagold-labeled RSSUFHC, detected by a gel shift, was imported and localized to the chloroplast stroma (Figs. 4A and 6A). The labeled mature protein was protease-protected and detected in the soluble stromal fraction (Fig. 4A). To control for the effect of protein modification on import, a smaller 12-14 Å particle, fluorescein maleimide, was conjugated to RSSUFHC and also imported into the chloroplast stroma. Another RuBisCO small subunit construct was made with a single internal cysteine at residue S58 (RSSU58CFH). Since the diameter of a linear protein chain is 4-6 Å, Undeccagold conjugated to the internal cysteine produces an effective probe diameter of at least 24 Å (25). The internally labeled Undeccagold probe was also imported into the chloroplast stroma, indicating the TOC/TIC pore size is greater than 24 Å (Fig. 4B).

**Fig. 4.**
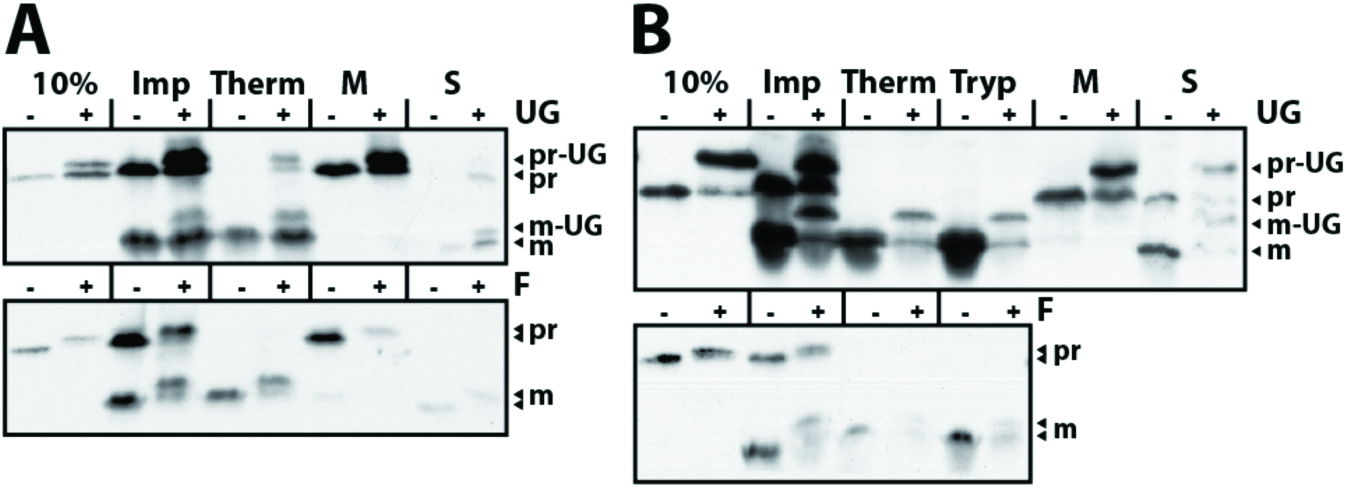
The TOC/TIC pore size is greater than 24 Å. Undeccagold (UG) and fluorescein (F) labeled RSSUFHC **(A)** or RSSU58CFH **(B)** were localized to the chloroplast stroma. Import reactions were conducted for 20 minutes and stopped in cold IB (Imp). For protease treatments, chloroplasts were further resuspended in 400 μg/mL thermolysin (Therm) or 240 μg/mL trypsin (Tryp) 30 minutes on ice and quenched with 12.5 mM EDTA or resuspended in 1 mg/mL SBTI. All samples were reisolated on 40% Percoll cushions and washed with IB. After reisolation, some non-protease treated samples were separated into membrane (M) and soluble (S) fractions by lysis in 2 mM EDTA for 10 minutes on ice and pelleting membranes at 16000 *xg.* Soluble fractions were precipitated in 15% TCA and washed with cold acetone. Samples were analyzed by SDS-PAGE and α-FLAG blotting.

**Fig. 5.**
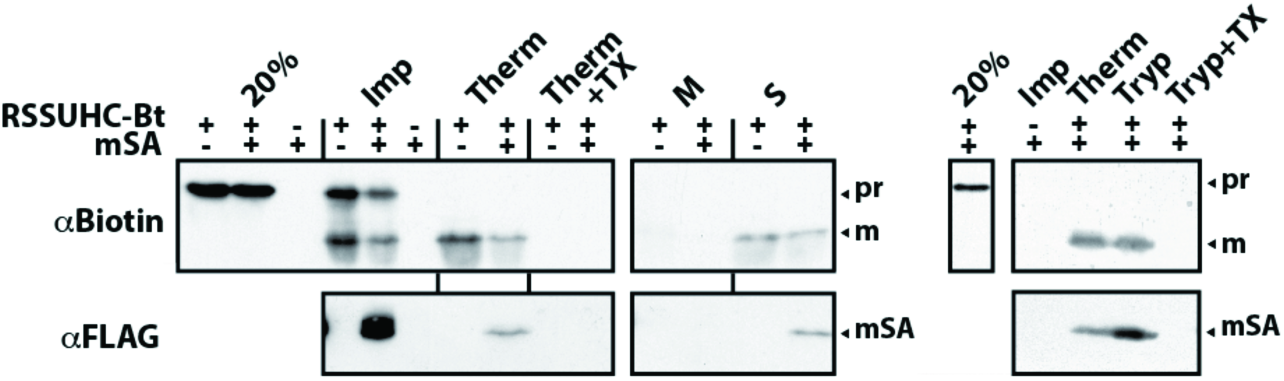
The TOC/TIC pore size is greater than 26.5 Å. Biotinylated RSSUHC (RSSUHC-Bt) was incubated with 4-fold molar excess mSA in IB prior to import reaction. Stromal localization of mSA was determined as in Fig. 4, except that the membrane/stroma fractionation was done on thermolysin treated chloroplasts and trypsin treatments were conducted at 25 °C. Additionally, protease treatments were conducted in the presence of 1% Triton X-100 (TX). RSSUHC-Bt and mSA were detected on α-biotin and α-FLAG blots, respectively.

**Fig. 6.**
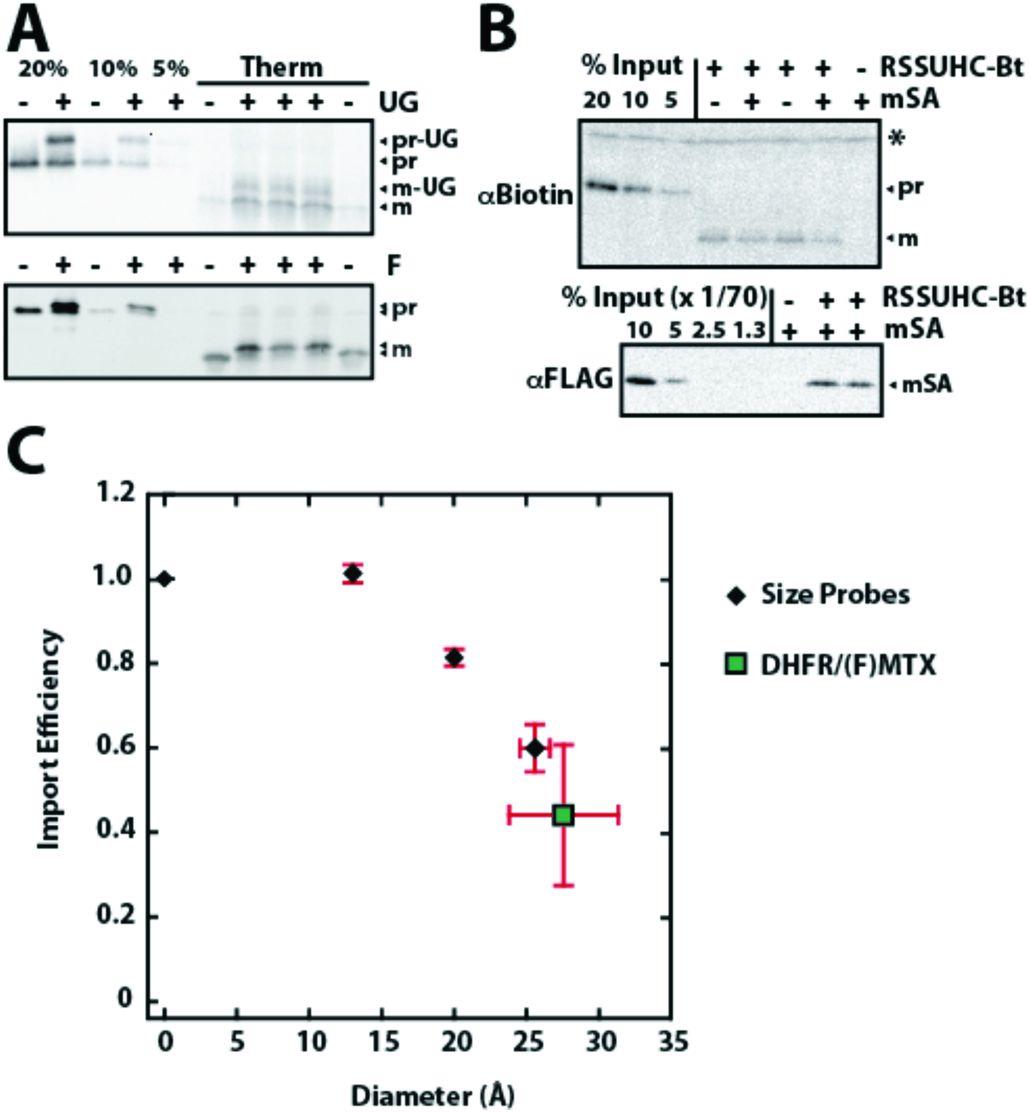
Import efficiencies of fixed-diameter probes. Import reactions of fluorescein (F) and Undeccagold (UG) labeled RSSUFHC **(A)** and mSA labeled RSSUHC-Bt **(B)** are quantified **(C).** RSSUHC-Bt (8.3 μg/mL in import reaction) was incubated with a 70-fold molar excess of mSA in IB prior to import. All reactions were stopped after 8 min with cold 400 μg/mL thermolysin and incubated 1 hour on ice. Samples were then treated as in Figs. 4 and 5. Quantifications were plotted as the labeled:unlabeled mature protein ratio. The internal control unlabeled protein was used for fluorescein and Undeccagold quantifications. Error bars (SD) for fluorescein, Undeccagold, and mSA import efficiency represent multiple replicates from 1, 2, and 2 independent experiments, respectively. tp22EcDHFR/FMTX import efficiency was plotted as the ratio of tp22EcDHFR import in the presence and absence of FMTX from Fig. 3F. Protein dimensions are given as minor axis diameters with error bars representing standard deviation away from a perfectly circular cross section in the minor axis plane. *nonspecific chloroplast protein.

The TOC/TIC translocons were additionally probed with cylindrical monomeric streptavidin (mSA) with a 25.6 Å diameter and a 2.8 nM K_d_ for biotin (26). His-tagged RuBisCO small subunit was biotinylated at a single C-terminal cysteine (RSSUHC-Bt), to which mSA was non-covalently bound. Although mSA is not a rigid particle like Undeccagold, it can still be used as a size probe since it must remain folded to retain affinity for RSSUHC-Bt during import. Without RSSUHC-Bt present, mSA did not bind to chloroplasts (Fig. 5). When imported with RSSUHC-Bt, mSA was localized to the chloroplast stroma, indicating the maximum TOC/TIC pore size is greater than 25.6 Å. Imported mSA was found in the soluble stromal fraction and was protease-protected from thermolysin and trypsin, but not in the presence of Triton. mSA import into chloroplasts was saturated above a 35-fold molar excess of mSA over RSSUHC-Btin the import reaction (Fig. S6). Under saturating conditions, mSA imported into chloroplasts at a 1:1 ratio with RSSUHC-Bt, ruling out the idea that the RSSUHC-Bt/mSA complex was dissociated by a strong unfoldase activity applied on mSA at the chloroplast membrane surface (Fig. 6B).

### The TOC/TIC Pore Size Is Larger Than That of TOM/TIM

The import efficiencies of the fixed-diameter probes were determined as the ratio of probe-labeled to-unlabeled precursor imported after eight-minute reactions. The eight-minute time point falls at the end of the linear portion of the reaction, and therefore correlates with the initial import rate (Fig. S6). As is to be expected, the probe import efficiency decreased with increasing probe diameters (Fig. 6C). Protein dimensions are given as minor axis diameters since the longest major axis can, in principle, be imported perpendicular to the TOC/TIC pore diameter. Thus, these minor axis values represent minimum pore size dimensions. The import efficiency of mSA supports the model of folded DHFR import since they have roughly similar minor axis diameters. The probes were imported more efficiently into chloroplasts than identical (20 Å Undeccagold) or similarly sized probes (26 Å Nanogold) were imported into mitoplasts through the TIM complex as determined by Schwartz and Matouschek (Fig. 6C)(27). In our hands, the Nanogold-precursor conjugate was unstable in the presence of chloroplasts and could not be used as a probe.

## DISSCUSION

In this study, we readdressed the tolerance of the TOC/TIC translocons for folded proteins and measured their functional pore size. We show that the DHFR/(F)MTX complex can be imported in a folded conformation by three independent methods: 1) By using subsaturating MTX concentrations, the possibility of independently imported MTX quantitatively reassociating with imported DHFR was ruled out. 2) FMTX import was significantly greater during concurrent DHFR import, indicating that DHFR/FMTX was imported as a complex. 3) The TOC/TIC pore size was greater than 25.6 Å, which is larger than the TOM/TIM pores and large enough to accommodate folded DHFR. Tetrameric avidin has previously been shown to block chloroplast import, suggesting the pore size has an upper limit of 50 Å (28). However, based on the reduced import efficiencies of mSA and DHFR/FMTX (Fig. 6C), the maximum pore size appears likely to be no more than 30-35 Å. In mitochondria, the TIM complex pore size is only slightly larger than 20 Å (27). The TOM pore is larger than the TIM pore, but less than 26 Å. Here, the TOC pore size was not determined independently of TIC, but it is tempting to speculate that the TOC pore may be larger since the inner membrane is solute selective, while the outer membrane is not (29). Since FMTX does have some inhibitory effect on DHFR import efficiency, DHFR is likely imported in a somewhat unfolded state in the absence of FMTX. The foldedness of proteins with diameters roughly 20-30 Å during import is probably determined by both the stability and size of the protein. Proteins below 20 Å are likely to remain folded during their import, while those above 30-35 Å are probably at least partially unfolded. However, given the pore size that we measured here, even large proteins may retain significant amounts of residual structure based on their particular mechanical unfolding pathways.

Prior evidence for protein foldedness during chloroplast import has mostly been limited to measuring protease sensitivity of tightly folded precursors bound to the chloroplast surface. These studies with different precursors yielded conflicting results. Purified ferredoxin reductase was found to be protease-resistant when bound to intact chloroplasts, but a purified chimeric OE33-RicinA protein was observed to be protease-sensitive (30, 31). An *in vitro*-translated ferredoxin-DHFR fusion protein complexed with MTX was found to be protease-sensitive when bound to purified envelope membranes (32). *In vitro*-translated plastocyanin-DHFR fusion protein complexed with MTX was found to be protease-sensitive when bound to intact chloroplasts (19). In this case, the protease may have cleaved the transit peptide from DHFR, leaving the resistant DHFR/MTX complex in the supernatant, which was not analyzed. The same study reported a lack of [^3^H]-MTX associated with chloroplast-bound DHFR, however, the [^3^H]-MTX was likely diluted away due to equilibrium debinding during chloroplast reisolation and washing, just as FMTX was debound by dilution in Fig. 3B. It seems unlikely that these tightly folded proteins could be unfolded by a passive mechanism at the chloroplast surface, yet an energy dependent unfoldase at the outer membrane has not been found. The quantitative import of mSA also indicates that there is no global unfoldase activity at the chloroplast surface (Fig. 6B).

The length of loosely structured N-terminal extensions (including transit peptides) on tightly folded proteins is critical for import efficiency into both mitochondria and chloroplasts as the transit peptide/presequence must reach internal chaperones that provide the energy for unfolding and/or translocation. Mitochondria require an approximately 80 residue N-terminal extension to span two membranes and reach mtHsp70 in the matrix (33). Here also, the chloroplast data is conflicting, with ferredoxin reductase (FNR) requiring roughly 80 residues for efficient import and titin requiring only 60 (34, 35). It was suggested that the unfoldase activity for titin resided in the intermembrane space (IMS) such that the transit peptide only had to cross one membrane to reach the chaperone. However, the average minor axis diameters of FNR and titin are 40 Å and 22 Å, respectively, which suggests that titin does not necessarily need to be unfolded prior to chloroplast import, unlike FNR. The force required for mechanical unfolding of titin by atomic force microscopy is roughly twice the DHFR/MTX unfolding force, which strongly suggests titin would also remain folded during chloroplast import since it is both smaller and more stable than DHFR/MTX (36, 37). IMS ATPase activity has been implicated in precursor translocation through TOC when uncoupled from TIC, but no IMS chaperone has been directly implicated in the coupled TOC/TIC import process to date (9, 38, 39). While it remains possible for IMS ATPase activity to play a role in precursor unfolding, the simpler explanation for the FNR/titin discrepancy would be related to the respective sizes of the proteins. Accordingly, folded proteins too large to pass through the TOC/TIC pore likely require the 80 residue N-terminal extension for efficient import, as in mitochondria.

Titin provides an ideal translocon-mediated protein-unfolding model because its mechanical unfolding pathway has been elucidated by atomic force microscopy. With mechanical force applied to the N and C termini of titin, the N-terminal β-strand A unravels first at approximately 100 pN, followed by the next β-strand A’ at 200 pN, after which the rest of the protein is easily unfolded at lower force (36). Stabilizing mutations in the A strand inhibit import efficiency of titin into mitochondria, while destabilizing mutations enhance import (40). The localized N-terminal stability of proteins is important for mitochondrial import because transient N-terminal unfolding is rate limiting. In chloroplasts, destabilizing strand A’ mutations, but not strand A mutations, affected import efficiency, and it was concluded that chloroplasts have a fundamentally different unfoldase mechanism, possibly involving an IMS chaperone (41). An alternate explanation is that strand A mutations did not affect chloroplast import because titin is small enough to be imported as a folded protein and unfolding of the A strand does not significantly reduce the minor axis diameter of the protein. Unlike strand A, destabilizing mutations in strand A’ and the core of titin likely allow the entire protein to unfold, thereby increasing chloroplast import efficiency. Strand A’ mutations probably did not affect mitochondrial import because mtHsp70 turnover and not protein unfolding was rate limiting (40). Due to the complication of folded titin import into chloroplasts, the mechanism of TOC/TIC-mediated protein unfolding remains unclear.

It is unknown which TOC/TIC components, if any, are specifically involved in precursor unfolding. One of the proposed stromal chaperones, Hsp93, has been ruled out by Kovacheva *et al.* (42). They show that DHFR import efficiency into Arabidopsis *hsp93-V/III-1* double mutant chloroplasts was not affected by MTX, while DHFR import into WT chloroplasts was affected. In light of folded DHFR/MTX import, their interpretation might still hold if DHFR is unfolded in the absence of MTX, which would provide the differential import efficiency readout based on protein stability required for interpreting their experiment.

The import of folded proteins has structural implications for the TOC and TIC complexes. The transport pathways of substrates utilizing β-barrel translocases such as mitochondrial Tom40 and bacterial FhaC pass through the central channel of the β-barrels, as determined by extensive cross-linking studies (43, 44). By analogy, the transport pathway of Toc75 substrates should also be through the central channel, but this has not been tested. Although the central channel is too small to accommodate folded DHFR, it has been speculated that the weakly interacting first and last β-strands of Toc75 could open (2). These Toc75 β-strands are structurally similar to those of its bacterial ortholog, BamA, and dissimilar from those of other β-barrel translocases, which have more tightly interacting terminal β-strands. The emerging model of BamA function in OMP biogenesis involves the terminal β-strands opening and pairing with newly inserted β-strands of substrate OMPs (45). In this model, once the substrate OMP is fully inserted into the membrane, its terminal β-strands would be paired with the terminal β-strands of BamA forming one large β-barrel. The substrate OMP would subsequently bud off from BamA and close its β-barrel structure. This mechanism requires BamA to be somewhat flexible and lends credibility to the prospect of the Toc75 β-barrel opening and potentially forming a large pore in cooperation with other TOC components. Pore flexibility is certainly not uncommon and is important even for SecYEG (46), where restricting the opening of the lateral gate caused transport inhibition of soluble proteins (25). The Toc159 membrane domain has been cross-linked to precursor proteins and may therefore cooperate with Toc75 to form part of the functional pore (47). Multiple Toc75 subunits within the TOC complex may also cooperate to form the pore. It is much harder to speculate on the nature of the TIC pore since there is very little structural information and continued controversy over the identity of the complex components. If multiple TIC complexes exist, they may also have different maximum pore sizes. Both TOC and TIC pores are likely to be somewhat flexible and expandable to accommodate folded proteins.

The import of folded proteins through the TOC/TIC translocon raises many questions for future investigation. Is the pore size of TOC larger than TIC? What is the mechanism of chaperone action that allows for folded protein translocation? Finally, the biological relevance for the large TOC/TIC pore size needs to be understood. It will be of interest to find native chloroplast proteins that are imported in a folded conformation, perhaps even when tightly bound to a cofactor.

## MATERIALS AND METHODS

### Plasmid Constructs

*P. sativum* RSSU cDNA was cloned into pET23a with 5′ NdeI and 3′ XhoI restriction sites. The three native cysteines were mutated (C-1S, C41V, and C112V) and a single C-terminal cysteine was inserted after the 6xHis tag by QuickChange PCR, yielding the plasmid pET23a-RSSUHC. A C-terminal FLAG tag was added 5′ of the His tag to yield pET23a-RSSUFHC. The RuBisCO construct with a single internal cysteine (pET23a-RSSU58CFH) was created by making a S58C mutation on pET23a-RSSUFHC and religating into pET23a with NdeI and XhoI sites to remove the C-terminal cysteine.

The pET23a-tp22DHFR plasmid was cloned by fusing the first 79 residues of *P. sativum* RSSU from the plasmid pET23a-tp22GFP (48) with mouse cytosolic DHFR DNA from the plasmid AtPC-DHFR (49) flanked by 5′ PstI and 3′ XhoI sites. The pET23a-tp22EcDHFR plasmid was made by replacing RSSU residues 80-180 in RSSUHC-pET23a with *E. coli* DHFR residues 2-159 flanked by two FLAG tags and 5′ BamHI/3′ XhoI sites. tp22EcDHFR (from *E. coli)* was made because full-length tp22DHFR (from mouse) could not be expressed in *E. coli.* EcDHFR without the tp22 fusion was cloned into pET23a with 5′ NdeI and 3 ′XhoI sites.

pRSET-mSA was a gift from Sheldon Park (Addgene plasmid # 39860)(26). It encodes mSA with 6xHis and FLAG tags for purification and detection.

### Protein Production and Labeling

Proteins were expressed in *E. coli* BL21 cells and purified under native (EcDHFR) or denaturing (other purified proteins) conditions with Ni-NTA Agarose according to the manufacturer (Qiagen). mSA was refolded by rapid dilution into a stirred PBS (137 mM NaCl, 2.7 mM KCl,10 mM Na_2_HPO_4_,1.8 mM KH_2_PO_4_ pH 7.2) solution overnight at 4 °C. Soluble mSA was concentrated and buffer exchanged into 50 mM NaCl, 50 mM HEPES pH 7.2 with Ultra-15 10K Centrifugal Filters (Amicon). EcDHFR and all other purified proteins were respectively buffer exchanged from elution buffer into 50 mM HEPES pH 7.2 or Labeling Buffer (8 M urea, 50 mM HEPES pH 7.2). Proteins were quantified on Coomassie-stained SDS-PAGE gels against BSA standards. [^3^H]-tp22DHFR was *in vitro*-transcribed and translated with T7 RNA polymerase and wheat germ extract, respectively, according to the manufacturer (Promega).

Maleimide reactions were carried out in Labeling Buffer containing 7 mM TCEP at room temperature for two hours. Fluorescein maleimide (Vector Labs), monomaleimido Undeccagold (Nanoprobes), and biotin maleimide (Sigma) stocks (10,1, and 100 mM, respectively) were made in DMSO. RSSUFHC or RSSU58CFH (1mg/mL) was reacted with 0.2 mM fluorescein maleimide or 0.05 mM Undeccagold and added directly to import reactions. RSSUHC was labeled with 5 mM biotin maleimide, precipitated with 15% TCA, washed with cold acetone, and resuspended in labeling buffer.

### Chloroplast Import Assays

Chloroplasts were isolated from 9-12 day old peas as described previously (50). Import reactions were conducted under 100 μE/m^2^s light with 3 mM ATP and isolated chloroplasts (0.33 mg/mL Chl) in Import Buffer (IB, 330 mM sorbitol, 3 mM MgCl_2_, 50 mM Tricine-KOH pH 8.0). Reactions with purified precursor protein contained 33 μg/mL protein unless otherwise specified. All protease treatments included 5 mM CaCl_2_. Those that included 1% Triton X-100 were subsequently precipitated in 15% TCA and washed with cold acetone. All samples were finally solubilized in 2X Laemmli Sample Buffer and analyzed by SDS-PAGE followed by fluorography or Western blotting. Input standards in Fig. 6B were diluted with cold Sample Buffer containing solubilized chloroplasts. Fluorography and fluorograph quantification were performed as previously described (48).

### Western Blotting

SDS-PAGE resolved proteins were transferred to PVDF membranes. α-FLAG blots were blocked with 5% fat-free milk in TBST (150 mM NaCl, 0.1% Tween 20, 50 mM Tris pH 7.5) and probed with monoclonal α-FLAG M2 (Sigma) diluted 1:1000 in blocking buffer. Goata-mouse IgG-HRP secondary antibody (SCBT) was diluted 1:10000 in blocking buffer. α-Biotin blots were blocked with 1% gelatin in TBST and probed with Avidin-HRP (Sigma) diluted 1:10000 in blocking buffer. Blots were developed with ECL substrate (GE Amersham) on film or on a ChemiDoc imager (Bio-Rad).

### Fluorescence Spectroscopy

Fluorescence measurements were conducted in a Fluorolog 3-22 spectrofluorometer (Horiba) set to λ_ex_=494 nm, λ_em_=518 nm, and 5 nm slit widths. Kinetics were measured in stirred 3 mL cuvettes.

## ACKNOWLEDGMENTS

We thank Dr. Jonathan Ho and Johnathan Keilman for cloning and purifying RSSUHC and tp22EcDHFR, respectively. We thank Drs. Kentaro Inoue, Hsou-min Li, and Enoch Baldwin for very helpful discussions and Anthony Ho for assistance with fluorimetry. This work was supported by NSF grant MCB-1330321 to SMT.

## SUPPLEMENTARY INFORMATION

**Fig. S1.**
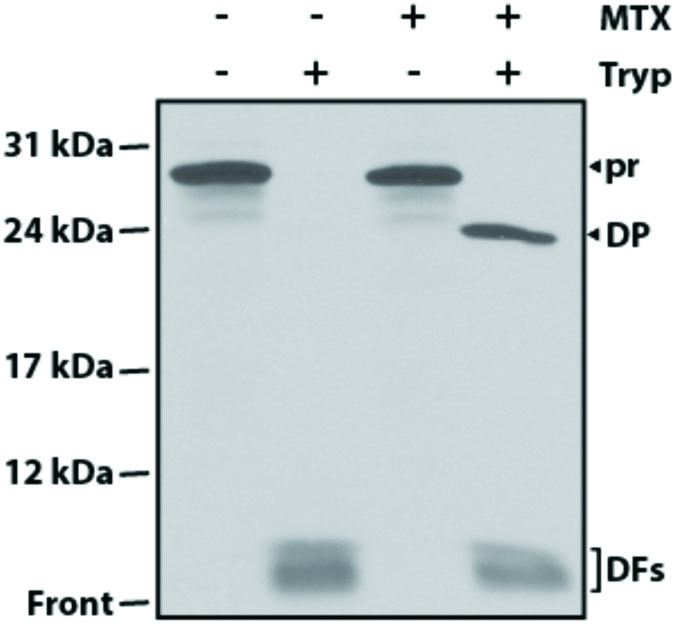
Stabilization of [^3^H]-tp22DHFR by MTX against trypsin digestion. *In vitro-*translated [^3^H]-tp22DHFR was incubated with 100 nM methotrexate (+, MTX) on ice for 10 min. Trypsin (+, Tryp) was then added to the above mixture reaching a final concentration of 200 μg/ml. Protease treatment was conducted at 37 °C for 5 min. Reactions were quenched with 2 mg/ml trypsin inhibitor (soy bean) on ice for 10 min, diluted in Sample Buffer and analyzed by SDS-PAGE and fluorography. pr, [^3^H]-tp22DHFR precursor; DP, DHFR degradation product; DFs, degradation fragments; Front, gel front.

**Fig. S2.**
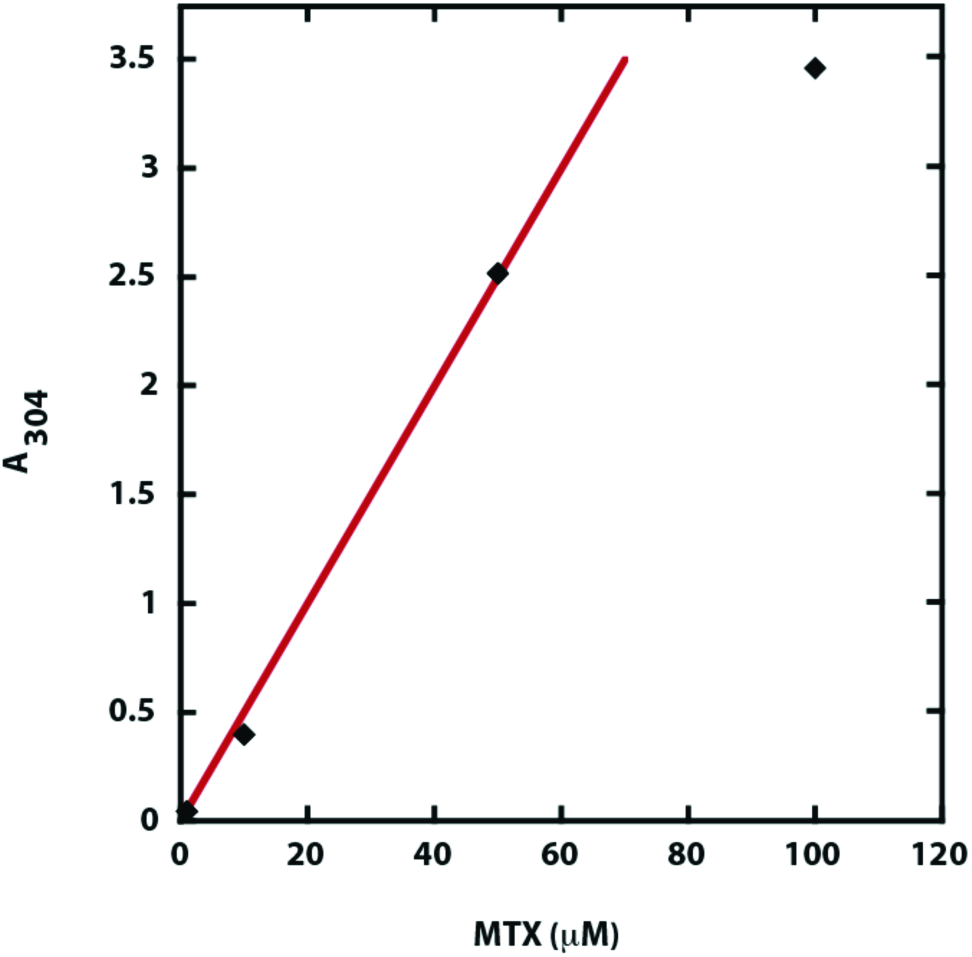
The A_304_ extinction coefficient of MTX in Import Buffer was determined to be 0.05 μM^−1^cm^−1^ within the linear range of 0 to 50 μM MTX. Within this linear range, the A_304_ of MTX decreased 33-fold after passing through a Zeba Spin Desalting Column (7K MWCO), which indicates a 660-fold decrease in MTX concentration.

**Fig. S3.**
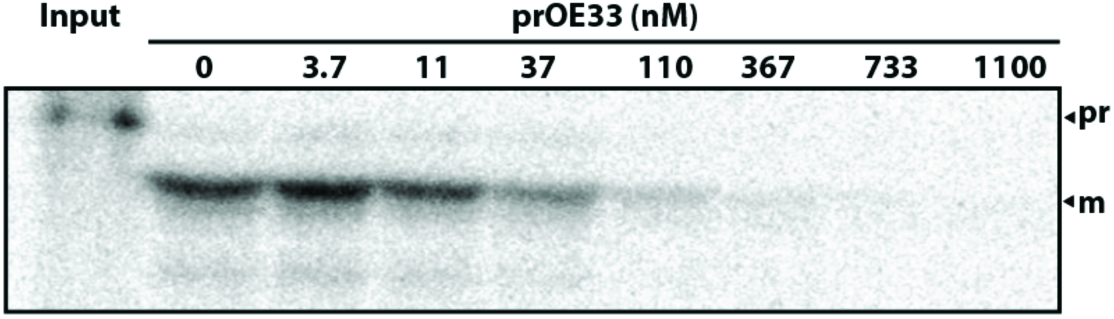
Import of [^3^H]-tp22DHFR competes with the precursor to the oxygen evolving complex 33 kDa subunit (prOE33) in the presence of MTX. His-tagged prOE33 was over-expressed in *E. coli*, purified under native conditions with Ni-NTA Agarose, and added at indicated concentrations to import reactions, while the concentrations of [^3^H]-tp22DHFR and MTX were held constant. Higher concentrations of prOE33 ultimately led to [^3^H]-tp22DHFR/MTX import quenching, indicating the native protein and the chimeric [^3^H]-tp22DHFR/MTX complex utilize the same TOC/TIC translocation machinery pr, [^3^H]-tp22DHFR precursor; m, mature [^3^H]-tp22DHFR.

**Fig. S4.**
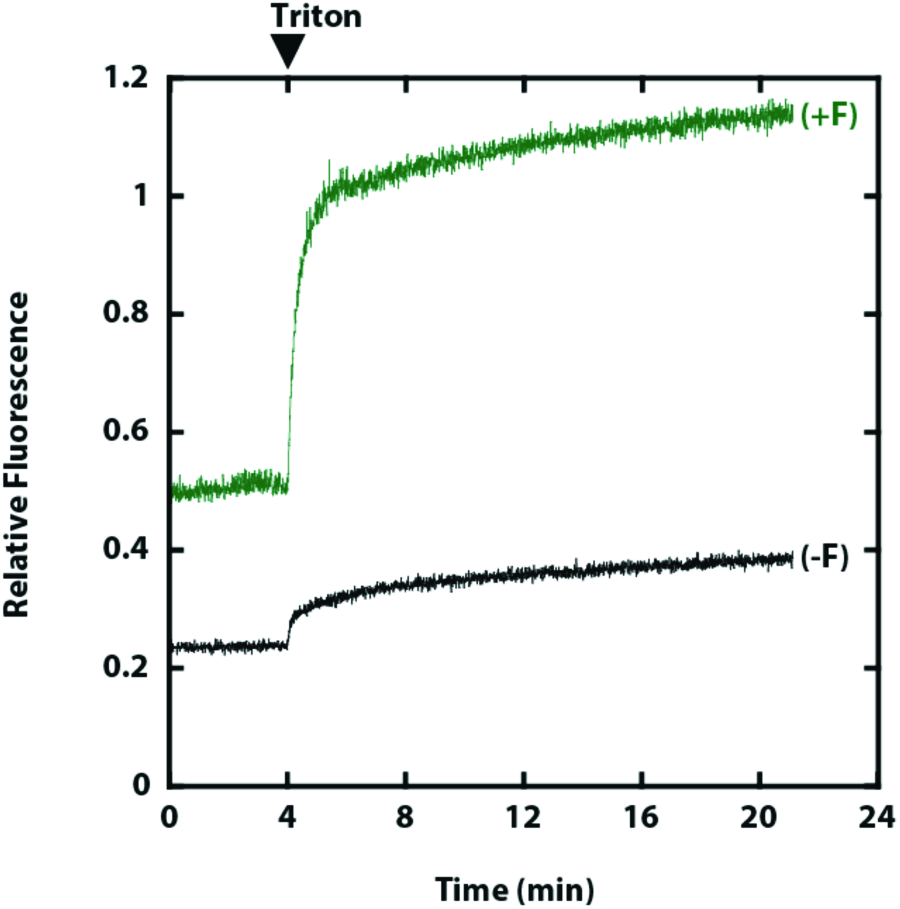
Fluorescence decay after chloroplast solubilization is specific to FMTX debinding and not fluorescein covalently conjugated to precursor protein. RSSUFHC labeled with fluorescein maleimide was imported into 0.33 mg/mL chlorophyll chloroplasts in a 240 μl reaction. Chloroplasts were thermolysin treated, reisolated, and assayed for fluorescence as in Fig. 3B. Triton X-100 was added as indicated. (+F), RSSUFHC-fluorescein imported chloroplasts; (-F), mock imported chloroplasts.

**Fig. S5.**
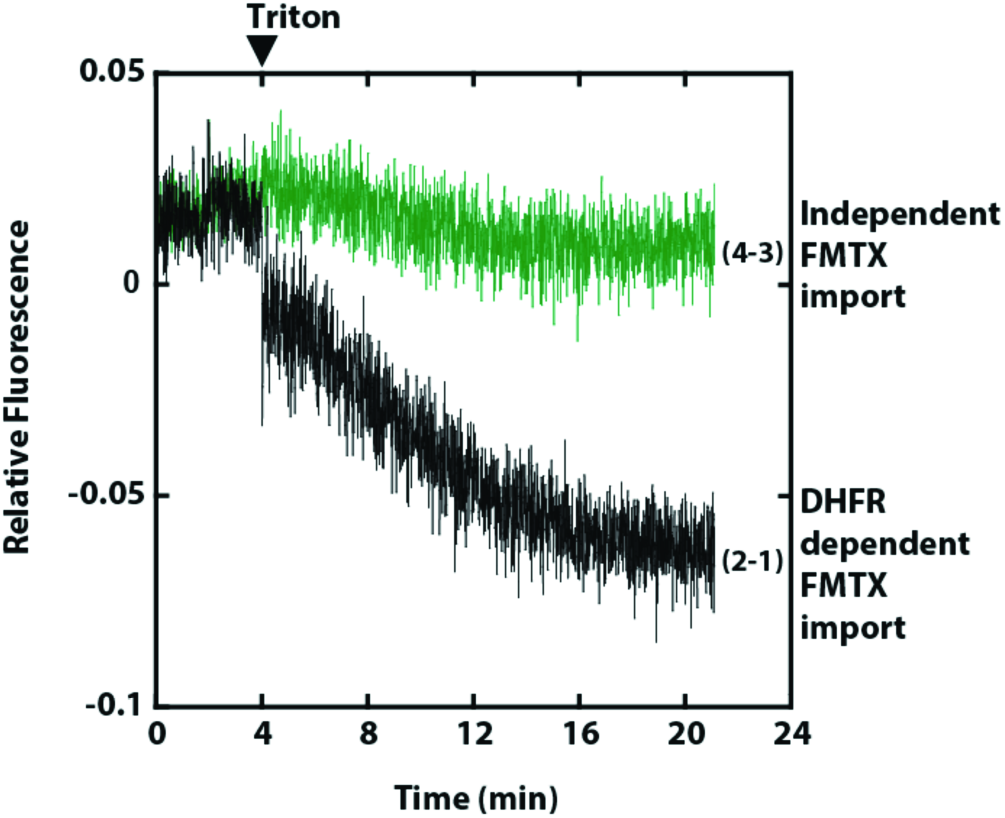
Kinetics of the MTX/FMTX binding replacement reaction. Derived from Fig. 3B, DMSO added traces were subtracted from MTX added traces (Traces 2-1 and 4-3)

**Fig. S6.**
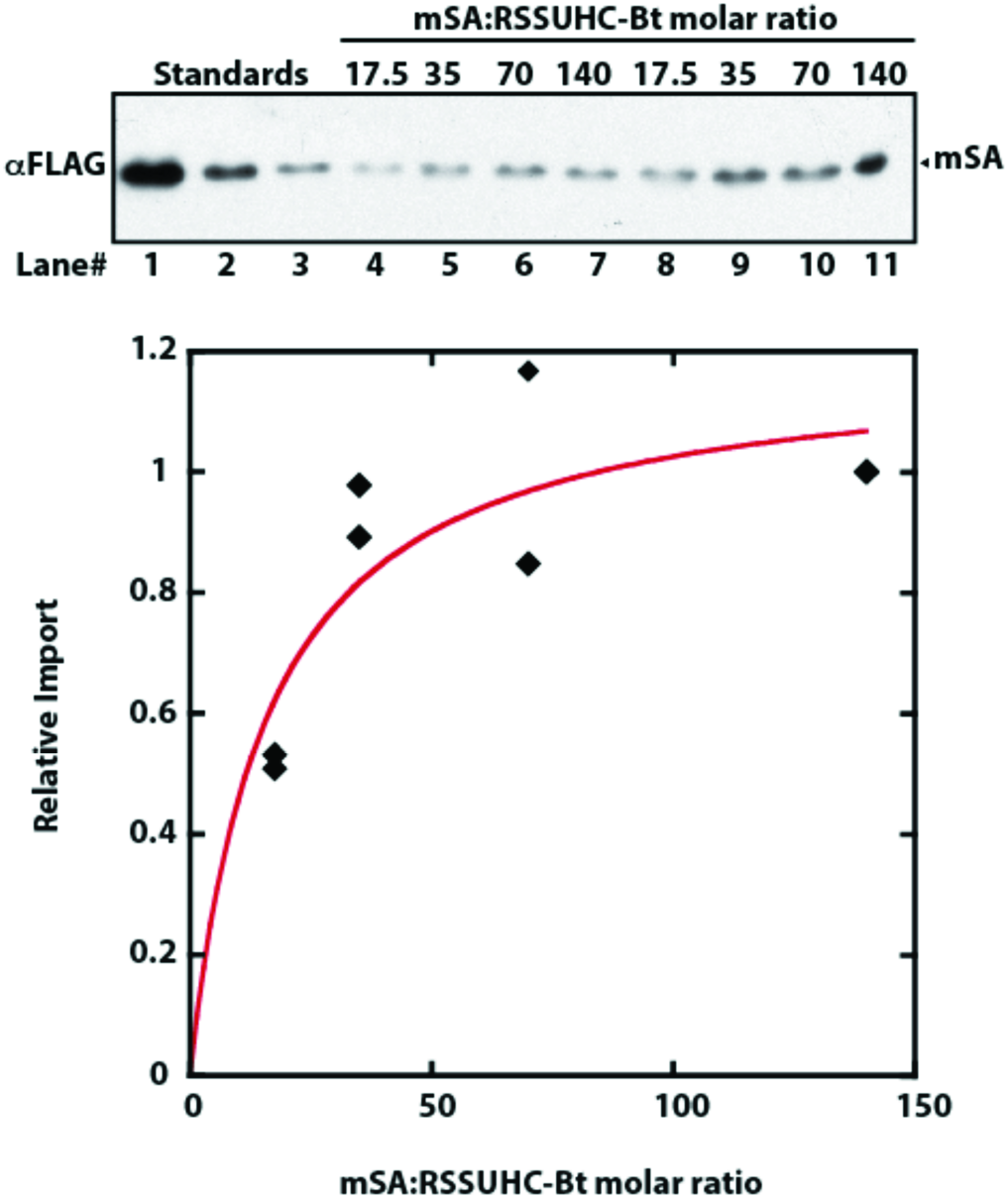
Saturation curve of mSA import. Import reactions were conducted as in Fig. 6B, except that chloroplast samples in lanes 8-11 were not reisolated on 40% Percoll cushions. RSSUHC-Bt was preincubated with excess mSA at indicated ratios. mSA import in lanes 4-7 and lanes 8-11 were each normalized to the 140-fold molar excess data points (lanes 7 and 11).

**Fig. S7.**
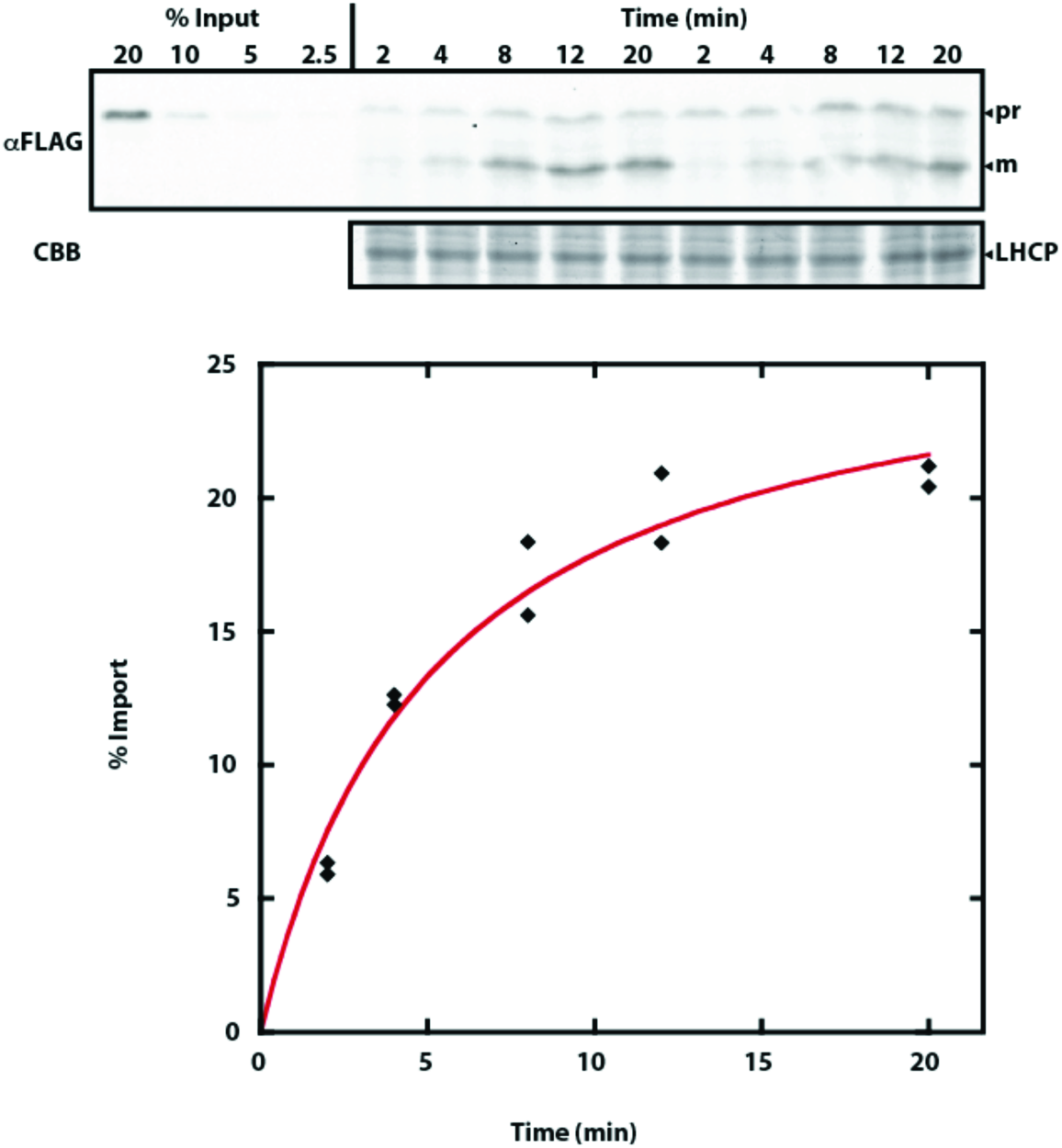
Time course import of RSSUFHC. Import reactions were stopped in cold 400 μg/mL thermolysin at indicated time points. Quantification of mature RSSUFHC is shown. CBB, Coomassie Brilliant Blue.

**Primer List:**

> Amplification of mouse DHFR with 5′ PstI and 3′ XhoI sites:
> F:CTGCAGATGGTTCGACCATTGAACTGCATCGTC
> R:CTCGAGTTAGTCTTTCTTCTCGTAGACTTCAAACTTATACTTGATG

> Amplification of *P. sativum* RSSU with 5NdeI and 3XhoI sites:
> F:ATCGCATATGGCTTCTATGATATCCTCTTCA
> R: GAATCTCGAGGTAGGATTCTGGTGTGTGGG

> RSSU C41V QuickChange PCR:
> F:GGAAGGGATGGGTTCCTGTCTTGGAATTTGAGTTGGAGAAAGG
> R:CCTTTCTCCAACTCAAATTCCAAGACAGGAACCCATCCCTTCC

> RSSU C112V QuickChange PCR:
> F:GACAACGTTCGTCAAGTTCAAGTCATCAGTTTCATTGCCCAC
> R:GTGGGCAATGAAACTGATGACTTGAACTTGACGAACGTTGTC

> RSSU C-1S QuickChange PCR:
> F:GGTGGAAGAGTAAAGAGCATGCAGGTGTGGCCTC
> R:GAGGCCACACCTGCATGCTCTTTACTCTTCCACC

> RSSUH-pET23a addition of C-terminal cysteine QuickChange PCR:
> F:GCACCACCACCACCACCACTGTTGAGATCCGGCTGCTAACAAAG
> R:CTTTGTTAGCAGCCGGATCTCAACAGTGGTGGTGGTGGTGGTGC

> Amplification of RSSU with 5Nde and 3XhoI and FLAG tag with 2xGly linker:
> F:ATCGCATATGGCTTCTATGATATCCTCTTCA
> R:GAATCTCGAGCTTGTCGTCATCGTCTTTGTAGTCTCCACCGTAGGATTCTGGTGTGTGGG

> RSSU S58C QuickChange PCR:
> F:CGTGAGCACAACAAGTGCCCAGGATACTATGATG
> R:CATCATAGTATCCTGGGCACTTGTTGTGCTCACG

> Insertion of BamHI and FLAG tag between residues 79 and 80 of RSSUFHC:
> F:AGTTGGATCCAGAGATCAGTTGTTGAAAGAAGTTGAA
> R:ACTCGGATCCCTTGTCGTCATCGTCTTTGTAGTCTGGTGGCAAATAGGAAAGAGTC

> Amplification of *E. coli* DHFR with 5′ BamHI and 3′Xho and FLAG tag with 2xGly linker:
> F:ATCTGGATCCATCAGTCTGATTGCGGCG
> R:GAATCTCGAGCTTGTCGTCATCGTCTTTGTAGTCTCCACCCCGCCGCTCCAGAAT

> Amplification of *E. coli* DHFR with 5′ NdeI and 3′Xho sites:
> F:ACTGCATATGATCAGTCTGATTGCGGC
> R: CTTACTCGAGCCGCCGCTCCAGAAT

